# Disentangling functional connectivity effects of age and expertise in long-term meditators

**DOI:** 10.1101/848507

**Authors:** Roberto Guidotti, Cosimo Del Gratta, Mauro Gianni Perrucci, Gian Luca Romani, Antonino Raffone

## Abstract

The effects of intensive meditation practices on the functional and structural organization of the human brain have been addressed by a growing number of neuroscientific studies. However, the different modulations of meditation expertise and of ageing, in the underlying brain areas and networks, have not yet been fully elucidated. These effects should be distinguished in order to clarify how long-term meditation can modulate the connectivity between brain areas. To address this issue, we tested whether meditation expertise and age can be predicted from the multivariate pattern of functional Magnetic Resonance Imaging connectivity, in Theravada Buddhist monks with long-term practice in two different meditation forms: Focused Attention (FA) and Open Monitoring (OM).

We found that functional connectivity patterns in both meditation forms can be used to predict expertise and age of long-term meditators. Our findings suggest that meditation expertise is associated with meditation-specific brain networks modulations, while age-related modifications are general and independent from the meditation type. Specifically, expertise modulated patterns during FA meditation include nodes and connections implicated in focusing, sustaining and monitoring attention, while the predictive patterns during OM meditation include nodes associated with cognitive and affective monitoring. Thus, the two forms of meditation may differentially contribute to counteract the effects of neurocognitive decline with ageing by neuroplasticity of brain networks.

## Introduction

Meditation can be characterized as a set of practices involving the regulation of attention, awareness and mental state (Lutz et al., 2008; Tang et al., 2015). Meditation practices can be usefully classified into two main styles – *focused attention* (FA) and *open monitoring* (OM) – depending on how attentional processes and cognitive monitoring are directed (Cahn and Polich, 2006; Lutz et al., 2008). In the FA style, attention is focused on a given object in a sustained manner, with related attentional regulation processes. The second style, OM meditation, is based on non-reactive monitoring of the contents of experience, primarily as a means to become aware of emotional and cognitive patterns.

Research suggests that FA and OM meditation styles involve different brain regions and processes, in association with their different phenomenological, attentional and cognitive features (Cahn and Polich, 2006; Lutz et al., 2015, 2008; Manna et al., 2010; Marzetti et al., 2014). Besides the involvement of brain regions in meditation, studies also emphasize the implication of brain networks associated with the regulation of attention, emotion and self (Malinowski, 2013; Tang et al., 2015). Indeed, new paradigms are emerging in cognitive, affective and social neurosciences that emphasize the interactive function of brain areas working together as large-scale intrinsically-connected brain networks (Fox et al., 2005; van den Heuvel et al., 2009). The study of brain networks and their interactions appears highly relevant for science of consciousness and meditation (Lutz et al., 2015; Malinowski, 2013; Raffone and Srinivasan, 2009). This approach provides new insights into how functionally connected systems in the brain, support or constrain, cognitive and affective functions (Bressler and Menon, 2010), also with implications for understanding attentional, monitoring and affective functions in meditation, and their differential involvement in FA and OM styles. Neuroscientific findings have also shown that the interactions within and between brain networks are altered in ageing (Hafkemeijer et al., 2013; Sala-Llonch et al., 2015). Other studies have shown the effectiveness of meditation training in transforming the dynamics of brain networks: remarkably, brain networks that are altered in ageing appear to be influenced through mindfulness practice in terms of enhanced cognitive, emotional and self-reference flexibility (Hasenkamp et al., 2012; Malinowski, 2013; Marzetti et al., 2014; Tang and Posner, 2014). FA and OM meditation forms may furthermore differentially contribute to modulate the activity within and between brain networks (Raffone et al., 2019).

Another important aspect in the neuroscientific research on meditation is given by the relationships between meditation expertise, age and neuroplasticity. Although this topic was addressed by several studies for both expertise (Brefczynski-Lewis et al., 2007; Lutz et al., 2009, 2008; Manna et al., 2010; Tomasino et al., 2012) and age (Kurth et al., 2017; Malinowski and Shalamanova, 2017), these investigations only compare different groups of subjects without directly assessing for the specific modulation of age and meditation expertise on the functional organization of the brain, including patterns of brain networks involved in the regulation of attention, emotion and self (Hölzel et al., 2011). It can generally be hypothesized that the efficiency of connectivity within and between such networks increases with meditation practice (expertise), whereas it decreases with ageing. Indeed, evidence suggests that meditation can contribute to maintain brain tissue and to preserve cognitive and emotional reserves (Kurth et al., 2017). It can moreover be hypothesized that FA and OM meditation expertise differentially affect functional connectivity patterns in the brain, in association with the attentional, monitoring and regulation processes involved in these two forms of meditation (Lutz et al., 2015, 2008). In particular, the recently developed Brain Theory of Meditation (BTM) (Raffone et al., 2019) suggests that FA meditation leads to a sharpening of activations and coupling between core brain networks, whereas OM meditation is characterized by a more distributed pattern of activations and higher coupling in particular in the left hemisphere. This theory suggests that these two forms of meditation enhance efficiency of neurocognitive processing in different ways, whereas neurocognitive ageing can be generally supposed to reduce efficiency of processing in the attentional, monitoring and regulation functions associated to both forms of meditation, besides a broad range of tasks.

A promising tool to address this issue is the study of the pattern of functional connectivity through a multivariate pattern analysis approach (MVPA). The interest in multivariate techniques in neuroimaging (Haxby, 2012; Haynes, 2015) has recently grown due to their ability to detect fine-grained differences in neurophysiological patterns as compared to univariate methods (O’Toole et al., 2007). Most MVPA studies focused on cognitive and perceptual aspects of brain functions (Cichy et al., 2014; Guidotti et al., 2015; Knops et al., 2009; Kragel and LaBar, 2016; Tosoni et al., 2016). More recently, with the increased availability of a large dataset, multivariate methods have been successfully used with functional connectivity, for the prediction of individual subject’s traits (Cole and Franke, 2017; Dosenbach et al., 2010; Finn et al., 2015; Woo et al., 2017).

MVPA has also been used in meditation studies to predict the age of a group of meditators using voxel-based morphometry (Luders et al., 2016) and to classify patterns of functional connectivity before and after a body-mind training course (Tang et al., 2017).

Here, to increase our understanding of the influences of meditation practice on the modulation of brain networks, we studied the modulation of a set of *brain networks* related to meditation expertise and age. Furthermore, we investigated the relationships between different forms of meditation and their effect on specific brain networks.

Given these aims, by using pattern regression, we sought to disentangle the modulation of meditation expertise and age on these networks. In particular, we assessed whether patterns of functional connectivity are predictive of the number of years of meditation practice (meditation expertise) in a group of long-term meditators, Theravada Buddhist monks, who practice meditation in a monastic context in which both FA and OM facets of meditation are emphasized. In addition, since the age of the participants was correlated with their expertise, we sought to predict age from the functional connectivity matrices. More specifically, we hypothesized that functional connections are differentially modified by ageing and meditation. Also, due to the specific cognitive demands of each meditation style, the connections that are modified by meditation were hypothesized to be different in FA and OM meditation styles, whereas those modified by age were hypothesized to be the same in the two meditation styles.

Therefore, according to the above remarks, we hypothesized that: *(i)* the patterns of functional connectivity within and between multiple functionally relevant brain networks can be used to predict years of meditation practice (expertise) as well as age, *(ii)* age modulated patterns are independent from the type of meditation, and *(iii)* expertise modulated patterns differ between FA and OM meditation forms. In particular, we hypothesized that the connectivity pattern modulated by meditation expertise in FA meditation includes nodes and connections implicated in focusing, sustaining, and monitoring attention and the connectivity pattern modulated by meditation expertise in OM meditation includes nodes and connections associated with cognitive and affective monitoring and control, as well as interoception (Lutz et al., 2008; Raffone et al., 2019).

## Materials and Methods

### Participants

Twelve Theravada Buddhist monks (males, mean age 37.9 years, SD 9.4 years) from Santacittarama Buddhist Monastery, in Central Italy, following a Thai Forest Tradition, participated in our study. Participants practiced FA (Samatha) and OM (Vipassana) meditation forms in a balanced way in this tradition, including silent meditation retreats (3 months per year). Meditation expertise is measured in years since the beginning of meditation practice in the monastic context (mean 16.41, SD 7.69 years). In this tradition, the monks typically practice Samatha–Vipassana meditation, with a balance of FA and OM meditation, 2 h per day with the monastery community, with a regular intensification of practice (with several meditation sittings during the 3 month Winter retreat. Thus meditation expertise can be measured by years of practice, with each year assumed to correspond to 1,200 hours of meditation practice with balanced FA and OM meditation facets, as suggested by the monks. The experiment was conducted with the subject written informed consent according to the Declaration of Helsinki, as well as with the approval of the Ethics Committee of “G. d’Annunzio” University of Chieti-Pescara.

### Experimental design

The experimental design consisted of three blocks of the following sequence: 6 min of FA and 6 min of OM meditation blocks intermixed with a 3 min non-meditative resting state block (Figure 1A) and cued by vocal instructions. The total duration of the experiment was 57 min. Instructions on the mediation type vs. rest were provided verbally before the beginning of each block.

**Figure 1.**
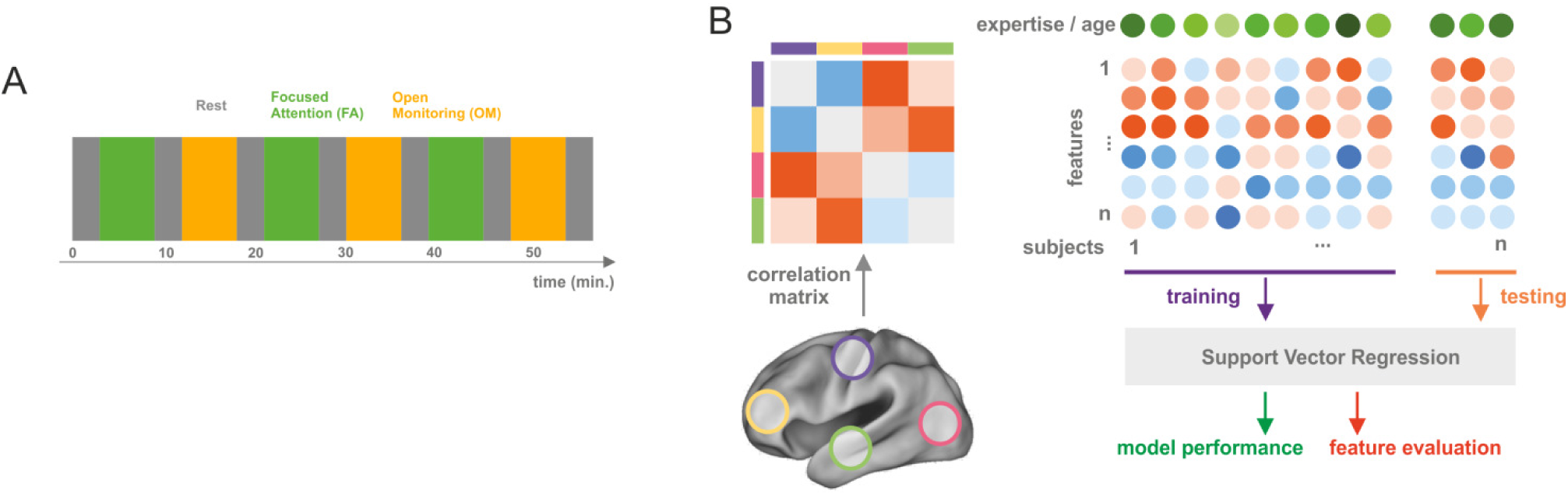
Experimental paradigm and analysis schema. Panel a) shows the experimental procedure consisted of three blocks of the following sequence: 6 min FA and 6 min OM meditation blocks intermixed with 3 min non-meditative resting state block. Panel b) shows the analysis schema consisted in extracting the average timecourse of the preprocessed BOLD signal (see Materials and Methods section) from 90 ROIs, and computing the pairwise Pearson correlation matrix between extracted timecourses; then the upper part of the correlation matrix was used as feature set for training a Support Vector Regression (SVR) model, to predict either the meditation expertise or the age of the participants; before training, a correlation based feature selection was performed to reduce the feature set. We split the dataset into two parts: 75% of the subjects were used for training and 25% repeating this procedure 50 times with permutation of the set of subjects for cross-validation. Finally, performance was evaluated using mean squared error (MSE) and correlation (COR), while feature evaluation was performed by extracting the selection frequency of the features and by inspecting the weights of the SVR model.

The following instruction were given for FA meditation: “gently engage in sustaining the focus of your attention on breath sensations, such as at the nostrils, noticing with acceptance and tolerance any arising distraction, as toward stimuli or thoughts, and return gently to focus attention on the breath sensations after having noticed the distraction source”. In OM meditation, participants were given the following instruction: “observe and recognize any experiential or mental content as it arises from moment to moment, without restrictions and judgment, including breath and body sensations, percepts of external stimuli, arising thoughts and feelings”. The instruction for Rest was the following: “rest in non-meditative a relaxed awake state”. Retrospective self-reports of the meditators suggested that they performed with accuracy both FA and OM forms during the experiment.

### Data acquisition

Data acquisition was performed on a 1.5 T Siemens Magnetom Vision Scanner, BOLD signal images were obtained using T2*-weighted echo planar (EPI) sequence with: TR = 4.087 s, 28 slices, voxel size 4 mm × 4 mm × 4 mm, 860 functional volumes.

A high-resolution T1-weighted whole-brain image was also acquired at the end of each session via a 3D-MPRAGE sequence (sagittal matrix = 256 × 256, FOV = 256 mm, slice thickness =1 mm, no gap, in-plane voxel size = 1 mm × 1 mm, flip angle = 12°, TR/TE = 9.7/4.0 ms).

### fMRI preprocessing

Preprocessing of raw images was carried out using Brain Voyager QX 1.7 software (Brain Innovation, The Netherlands). The first five scans were discarded to reach T1 saturation. Preprocessed functional volumes were co-registered with the corresponding structural data set. The co-registration transformation was determined using the slice position parameters of the functional images and the position parameters of the structural volume. Temporal filtering included linear and non-linear (high-pass filter of two cycles per time course) trend removal. No spatial or temporal smoothing was applied.

### Regions of Interest (ROI)

We selected the functional ROIs parcellation defined by Shirer et al. (Shirer et al., 2012), which is set of ROIs that broadly covers a large part of the cerebral cortex using a functional subdivision. The authors extracted 90 ROIs from 14 networks obtained with ICA from resting state fMRI data: visuospatial (VSN), high (hVN), and primary visual (pVN), left (LECN), and right executive control (RECN), posterior (pSN), and anterior salience (aSN), dorsal (dDMN), and ventral default mode (vDMN), sensorimotor (SMN), basal ganglia (BGN), language (LanN), auditory (AudN), and precuneus (PCuN) networks. The selected ROI list is presented in Table S1.

### Preprocessing for the functional connectivity analysis

We segmented anatomical images into grey-matter, white-matter, and cerebrospinal fluid (CSF) using the FSL FAST algorithm (Zhang et al., 2001) to extract masks for the subsequent preprocessing steps. White matter and CSF average signals obtained from segmented masks were regressed out from the fMRI signal, together with movement parameters. Then, the residual signal was filtered using band-pass filter (0.009 – 0.08 Hz), and finally scrubbed (Power et al., 2012) with a FD equal to 0.5, the resulting signal was used for all the subsequent analyses. Since ROI coordinates were given in the MNI space, a non-linear co-registration from the participant brain space to the MNI template was performed, and then the inverse of the obtained transformation was applied to the defined ROIs in order to extract the signal from the subject space. After the inverse transformation was computed, the intersection mask between the grey matter mask and all the defined ROIs (see Region of Interest) was found in the subject space. Due to uncertainties in this process a few small size ROIs in some participants had no intersection with grey-matter. We therefore selected only the ROIs that had an intersection with grey matter in all participants. Overall, only 7 ROIs were discarded from a total of 90 ROIs.

### Functional connectivity analysis

The average ROI time course was extracted from each ROI and the Pearson’s correlation coefficient between time courses was computed between all the ROIs obtained for each subject, independently from the meditation modality, and for each experimental block. Then, Fisher’s z-transformation was applied to ensure normality, and thus a square correlation matrix for each block was obtained.

### Pattern Regression

The correlation values from the upper (or lower) triangle of the correlation matrix were used as features for the regression (Fig 1B). Specifically, we computed correlation matrices for each participant, block, and meditation style. Therefore, the dataset for pattern regression analysis was composed of six correlation matrices per participant, resulting in 3403 features for 36 samples.

A correlation based feature selection (Dosenbach et al., 2010) was applied to reduce the dataset dimensionality, as the ratio between the number of samples and features was very large. We computed Pearson’s correlation between each feature and meditation expertise; then we chose the 2% features with the highest correlation as input to the regression algorithm.

We performed model selection by means of cross-validation. The dataset was split into two parts: 75% of subjects were used as a training set and the remaining 25% as testing set. To ensure independence of data, we used in the training dataset only data from a set of subjects, and in the testing dataset, only data from the remaining set of subjects. This procedure was repeated 50 times (Varoquaux et al., 2016), by shuffling subjects in the training set to provide a good estimator of the prediction error. Feature selection was applied at each repetition, and only to the training set, in order to avoid biasing of the prediction error (Pereira et al., 2009).

We analyzed the dataset using a linear Support Vector Regression (SVR), with *C* = 1. We then computed both the mean squared error (MSE) and the Pearson’s correlation (COR) between predicted and real values to validate the model. All these analyses were carried out with *scikit-learn* (Pedregosa et al., 2011) and *scipy/numpy* packages (Oliphant, 2007).

### Relevant feature analysis

We extracted the weights of each connection that were tuned during the training phase. Since we selected only correlated features before training the prediction model, we discarded noisy features that did not carry information about the target variable and that could thus bias the interpretation of the model feature weights (Haufe et al., 2014), therefore the weights of the model indicate the importance of each connection for the prediction task.

The weights were finally normalized to have a mean equal to 0 and a standard deviation of 1 and averaged across cross-validation folds, in order to obtain a single matrix with the contribution of each connection.

## Control analyses

Since the two variables we want to predict (age, expertise) can be correlated, they may represent a reciprocal potential confound for the prediction task. The main approaches to control confounds are: i) using the confound as a feature to check its predictive power, and ii) regressing out the confound from the data. These methods have important flaws that may bias the results (Snoek et al., 2018). The first method may effectively control the confound, but a good classification performance could still be misinterpreted due to noise of the confound (Haufe et al., 2014). The second removes the variance related to the confound, but may also remove some variance related to the variable of interest; moreover, in some cases it has been observed that regressing out the confound variable gave the same results as in the uncontrolled case (Rao et al., 2017).

Due to the above pitfalls, we trained the two separate models (Bron et al., 2015), one for the age, and one for the expertise, and we compared the features selected by the two models. If feature sets relevant for prediction overlap in the two models, then the prediction is biased, while if the relevant feature sets are disjoint, the prediction is not affected by the confound variable. This approach has many advantages: first, the target variable is directly predicted from the functional connectivity matrix; second, using a correlation-based feature selection we only include variables that share variance with the target variable, without removing the signal from a correlated confound as in regression. Finally, a post-hoc check for the confounding effect can be evaluated as the models are trained in the same way. A control analysis regressing out the confound from data has been carried out, and results are presented in Supplementary Material.

We did not perform any analysis using a control group, since on one hand, our analysis aims to predict the expertise of long-term meditators in two different meditation styles, thus we cannot predict years of experience in a non-expert group of subjects, and, on the other hand, the prediction of age from connectivity data acquired during meditation is difficult to compare since it may be biased, first, by the different meditation practice operated by the novices, and second, by the different life-styles between these groups, that would act as confounds in the comparison.

### Statistical analysis

The statistical significance of the regression results was assessed through permutation tests (Nichols and Holmes, 2002). In particular, we shuffled a thousand times (*n*=1,000) the prediction labels (i.e. years of meditation practice, and age) and repeated the entire prediction process, including feature selection and cross-validation, to obtain the null distribution for both MSE and COR errors. The reported p-value is the ratio between the number of permutations that outperform the case with no permuted labels and the number of permutations. The p-value is the probability of observing the reported error by chance using the null distribution obtained after permutations. Bonferroni correction was used to assess for Multiple comparisons.

## Results

### Prediction of years of meditation expertise

The prediction performance was calculated using the mean squared error (MSE) and the correlation coefficient (COR); statistical significance was assed using permutation tests with n=1000. We predicted the years of meditation expertise with a MSE of 0.497 (p<0.01; permutation test) and a COR of 0.857 (p<0.001; permutation test) in FA meditation. During OM meditation we predicted the expertise with a MSE of 0.520 (p<0.01) and a COR of 0.834 (p<0.001). The model average error of the model was about 0.5 years^2^.

### Prediction of age

Since the age of the participants may represent a potential confound, as it is correlated (r = 0.68) with the years of meditation practice, we trained a separate model to predict the age of the participants. The age of the meditators can be predicted with a MSE of 0.186 (p<0.005; permutation test) and a COR of 0.863 (p<0.001; permutation test) during the FA meditation, while during OM the MSE was of 0.201 (p<0.005) and the COR is of (0.791; p<0.001). The average model error was about 0.20 years^2^ for FA and OM.

### Feature selection frequency

We used a feature selection algorithm to reduce the number of connections to be used for the regression. We computed Pearson’s absolute correlation between each feature and the expertise (or the age) and then we chose the 2% features with the highest correlation as input to the regression algorithm. We performed the feature selection only in the training set in order to avoid the introduction of a bias in the prediction performance within a random cross-validation repeated 50 times. In this framework, the feature selection frequency is important to understand the importance of the connections for the predictive model.

We plotted the number of times a feature was selected by the feature selection algorithm for each cross-validation fold in Figures 2 and 3. Each dot represents a connection (feature) and the position coordinates *x, y* represent the number of times this connection was selected in the regression, in all cross-validation folds, for one of two different target variables (age and expertise respectively) (in Fig. 2), and in FA and OM meditation styles respectively (in Fig 3).

**Figure 2:**
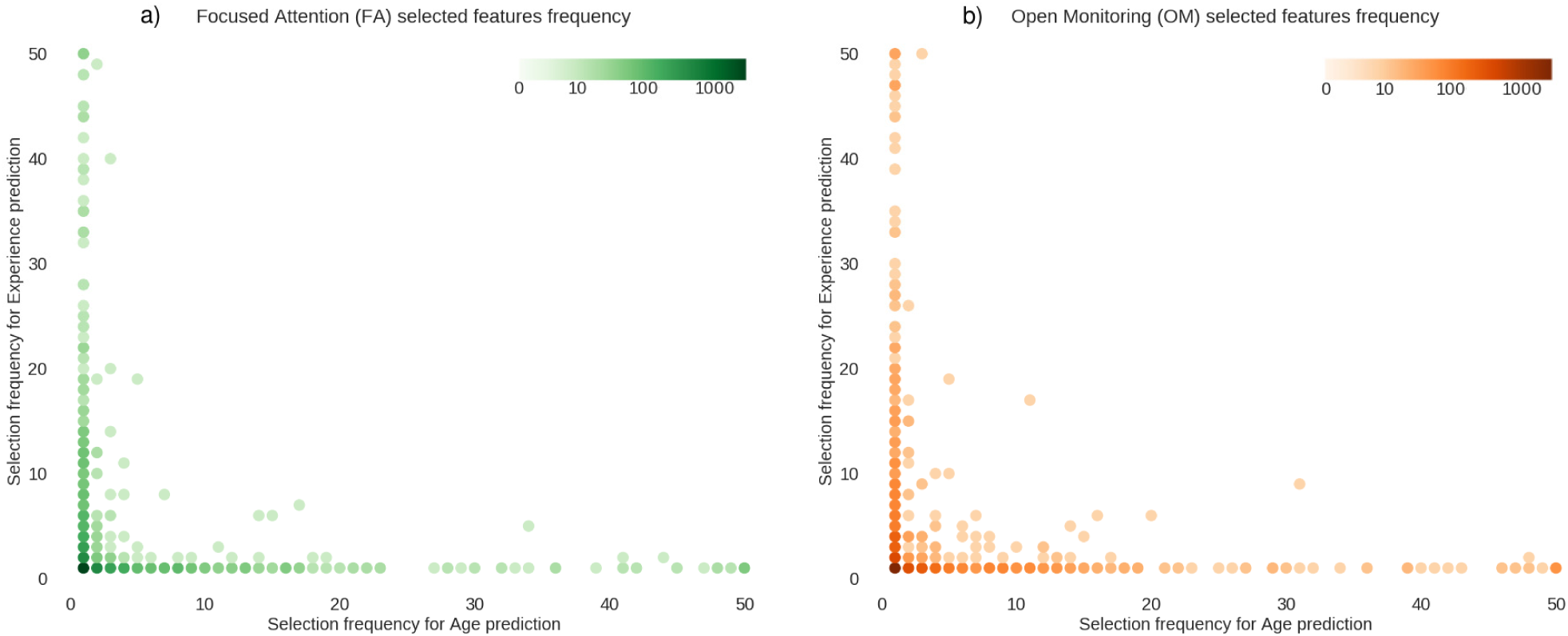
Feature selection frequency in FA and OM meditation styles. Figure shows the feature selection frequency for focused attention (panel a) and open monitoring (panel b) meditation, for age prediction (x-axis) and expertise prediction (y-axis). The maximum frequency value is equal to the number of cross-validation folds. The color of the dots indicates the number of features with that particular selection frequency.

**Figure 3:**
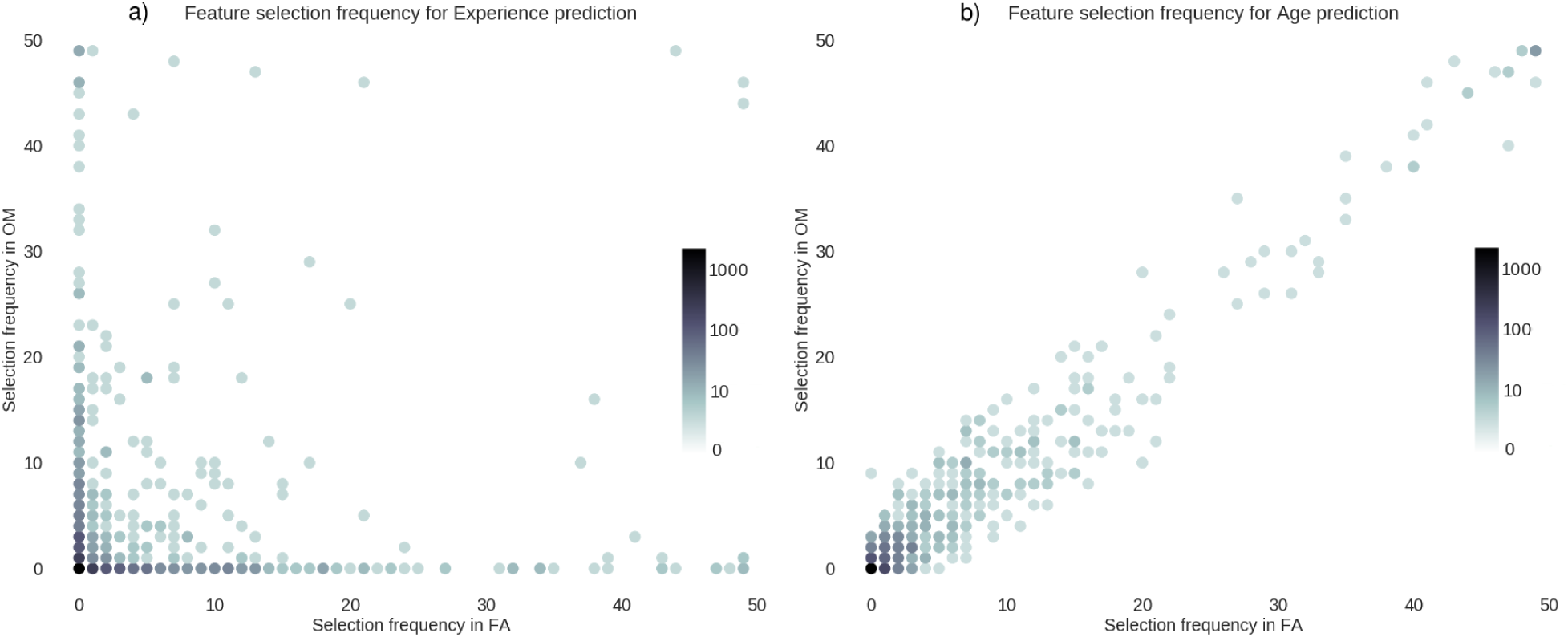
Feature selection frequency for expertise and age prediction. Figure shows feature selection frequency for meditation expertise prediction (panel a) and age prediction (panel b), for focused attention (x-axis) and open monitoring meditation styles (y-axis). The maximum frequency value is equal to the number of cross-validation folds. The color of the dots indicates the number of features with that particular selection frequency.

Figure 2 shows the sets of connections used to predict age and meditation expertise, in FA (Fig. 2a) and in OM meditation (Fig. 2b), respectively. In these plots, dots representing features with similar frequencies in the two meditation styles lie close to the diagonal, while dots representing features with different frequencies lie far from the diagonal. The figure suggests that the selection frequency of the largely correlated features is uncorrelated for the two prediction tasks. Indeed, the calculation of frequency correlation across features yielded a very low value (FA: *r*=-0.05, p>0.05; OM: *r*=-0.04, p>0.05). These results demonstrate that connections that are involved in age prediction are different from those involved in predicting the meditation expertise.

In Figure 3 we plotted the frequency of the selected features for prediction of meditation expertise (fig. 3a), and age (fig. 3b), respectively. Crucially, when features are used to predict the number of years of meditation practice (fig. 3a), feature selection frequencies appear to largely differ in the two forms of meditation (i.e. the dots lie mostly far from the diagonal). In other words, the two meditation styles recruit different connections, and this is confirmed by the low value (*r* = 0.14, p<0.05) of the frequency correlation across features. Instead, when features are used to predict the age of the meditators (Fig 3b), the algorithm selected the same features in the two meditation styles, and indeed the frequencies are highly correlated across features (*r* = 0.97, p<0.001). This further demonstrates that the connections affected by aging are distinct from those involved in meditation, as they appear to be insensitive to meditation style.

### Prediction-relevant feature weights analysis

Next, we analyzed the feature weights of the regression model to evaluate which nodes and functional connections were more predictive of meditation expertise and age in FA and OM meditation, and to what extent. As mentioned above, we trained 50 regression models each one for a different cross-validation split; therefore, 50 different sets of connections were used. We then restricted the analysis on those connections that were selected at least in 95% of the trained models (48 models). For each connection, the weights were averaged across cross-validated folds, and scaled with z-score.

We further plotted the ten most important nodes for the prediction of meditation expertise in FA and OM meditation (Fig 4a-b), and for the prediction of age in OM and FA meditation forms (Fig 4c-d). Values shown in Figure 4 are normalized absolute weights.

**Figure 4:**
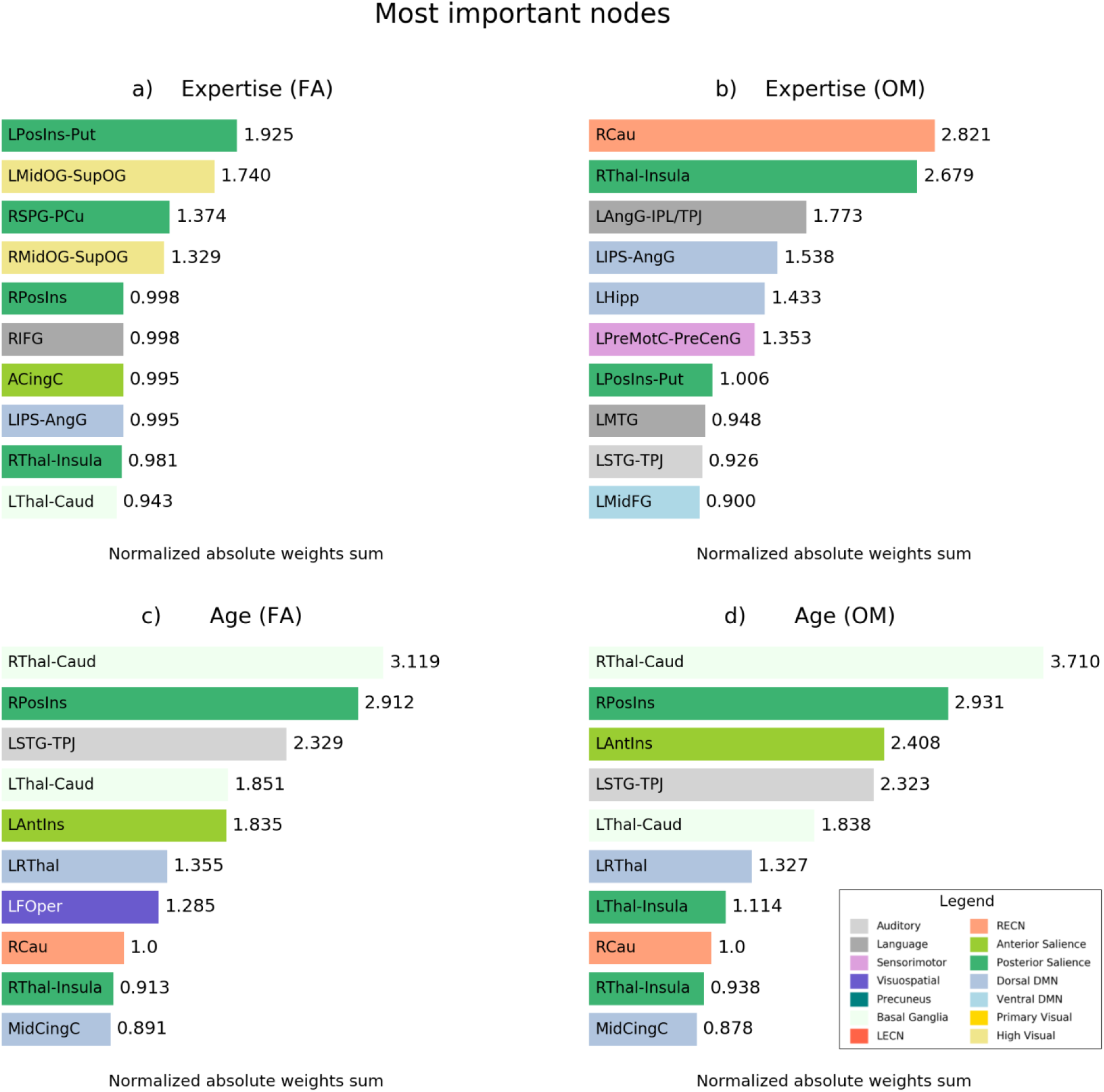
Normalized absolute weights of the nodes for different prediction task. Figure shows normalized absolute weights for expertise prediction in FA meditation style (panel a), in OM style (panel c), for age prediction in FA style (panel c) and in OM style (panel d). These weights are obtained by extracting the absolute weights learned by the model and averaging across-folds, then normalize them to have mean equal to 1 and standard deviation equal to 0. Finally, for each node we averaged the weights between connections that include that particular node.

As predicted from the results of the previous analyses, the most important nodes used to predict participants’ age were not only almost the same (9 out of 10) but they were also positioned in a similar order of importance in both meditation styles (panels c and d in fig. 4). Instead, the prediction of meditation expertise used different sets of nodes in the two meditation forms, with only two common nodes in the first ten (panels a and b in fig. 4), but with different rankings.

In Figure 5 we plotted the contribution of the connections in the prediction of meditation expertise in FA (fig. 5a), in OM (fig. 5b), and across both OM and FA meditation forms (fig. 5c). For the latter connections, we chose those that were selected 95% of the times together in OM and FA regression models.

**Figure 5:**
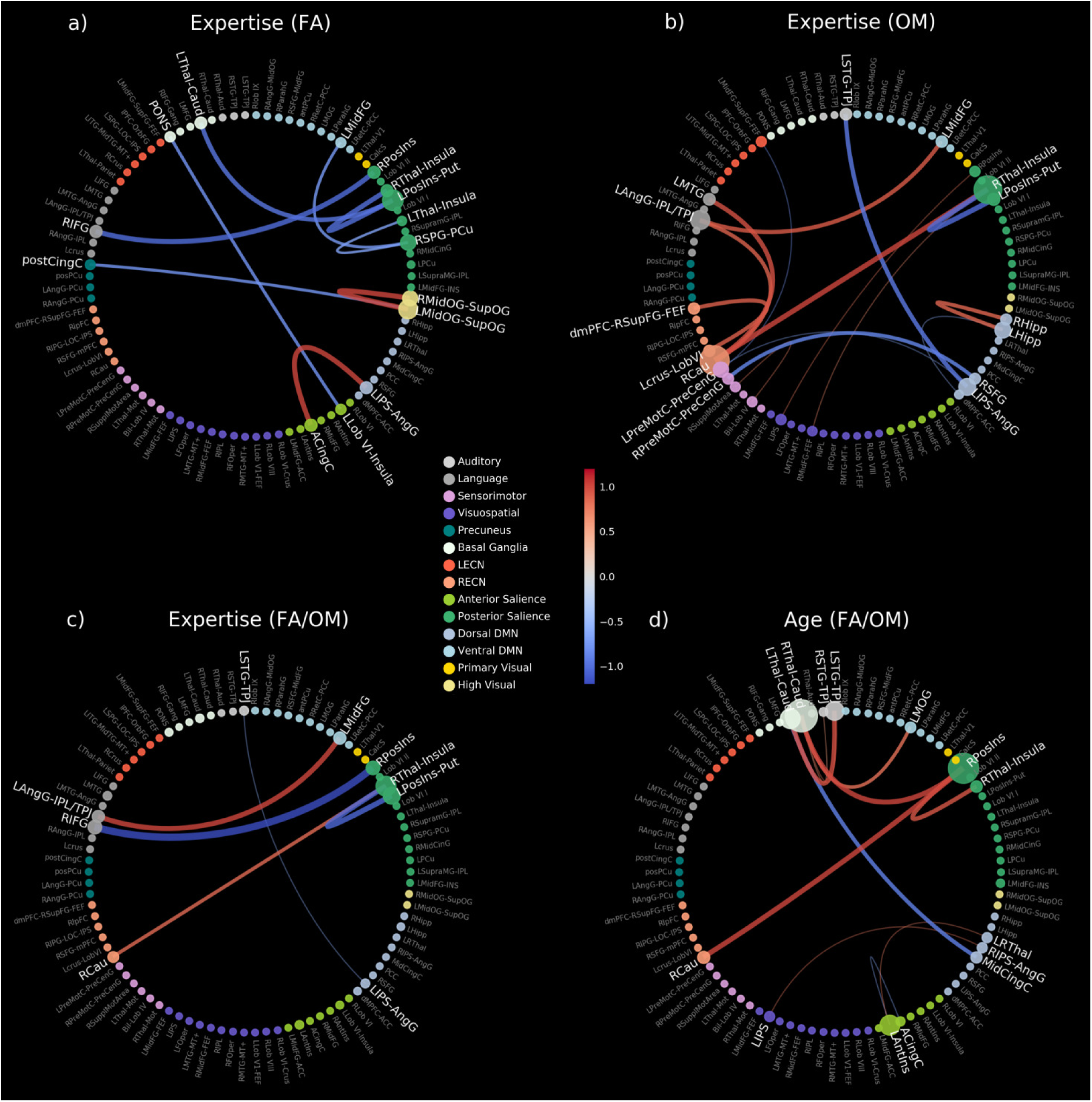
Normalized feature weights averaged across cross-validation folds. Weights used for the prediction of age (panel d) and meditation expertise (panels a, b, c) are shown.

## Discussion

We showed that meditation expertise and age can be predicted with good accuracy using fMRI functional connectivity from data recorded during focused attention (FA) and open monitoring (OM) meditation. Our findings demonstrate that patterns of functional connectivity are able to significantly predict years of meditation practice as well as age.

Our results also highlight a long term effect of the practice of meditation on functional connectivity and shed light on the different contributions of age and expertise on brain functional connectivity during meditation.

We found that age-modulated patterns are independent from the type of meditation while expertise modulated patterns differ between FA and OM meditation forms. Moreover, the two meditation styles we investigated were found to impact connectivity in different ways, and the connections they affected were consistent with their specific cognitive demands.

On one hand, the stability of the connectivity patterns modulated by age, across different meditation styles, suggests that the performed meditation moderately reorganizes age-specific connections, providing further evidence to the stability of brain networks (Gratton et al., 2018).

On the other hand, the meditation-specific modulation of brain networks due to expertise, demonstrates that long-term histories of co-activation among meditation-specific regions strongly impact the organization of brain networks (Gratton et al., 2018), supporting the idea that connectivity profiles could predict cognitive behavior (Finn et al., 2015).

In line with the Brain Theory of Meditation (Raffone et al., 2019), at a global level we observed a higher involvement of decreased functional connections with expertise in FA meditation, thus suggesting a sharpening of brain responses with FA meditation, plausibly linked to a more efficient allocation of brain resources to response-relevant stimuli in a given task (Manna et al., 2010). OM meditation was instead characterized by a more prominent increase of connections with expertise, with particular reference to connections within the left hemisphere and between the two hemispheres, also in line with such theory.

The meditation model of FA style proposed by Hasenkamp et al. (Hasenkamp and Barsalou, 2012) entails four stages: i) sustaining the focus of attention on the intended objects; ii) mind-wandering, iii) awareness of mind-wandering and iv) reorientation of the attention to the intended object.

Indeed, in FA meditation we found an increase in connectivity associated with meditation expertise between the left intraparietal sulcus – angular gyrus (LIPS-AngG) and the anterior cingulate cortex (ACC), two brain regions that are involved in top-down attention and attentional control.. In particular, ACC is crucial for monitoring conflicts (Botvinick et al., 2001; Petersen and Posner, 2012), such as the conflict between focus on the meditation object, and distractions, while IPS and AngG are directly involved in top-down attentional selection, and sustained attention (Thakral and Slotnick, 2009), two other important facets of FA meditation.

In addition, the white matter fractional anisotropy of ACC has been demonstrated to increase in short term meditators, this may be due to an increased communication efficiency and an improvement of attention monitor skills (Tang et al., 2012). The increase of the correlation of the ACC with IPS/AngG, as the expertise increases, may reflect an efficient communication process for detecting distractions and reorienting the attention as explained in the FA meditation model proposed by Hasenkamp and collaborators (Hasenkamp et al., 2012). We have also observed that meditation is associated with the formation of a series of connections with a negative correlation with meditation expertise, including the connection between posterior cingulate cortex (PCingC) and the left occipital gyrus (LmidgOG-LsupOG). This evidence can be explained in terms of a reduced occurrence of spontaneous thoughts and mental images during sustained attention in FA meditation. This is consistent with a series of results reported in the literature (Brewer and Garrison, 2014; Marzetti et al., 2014), and with the well established involvement of PCingC in mind wandering and stimulus-independent thought within the default mode network (Greicius et al., 2003; Pagnoni, 2012).

We also found that meditation expertise increase is associated with an increased connectivity within the higher visual network. This trend may be associated with a spontaneous focused visualization, though with closed eyes, called *nimitta* in the Buddhist meditation tradition that may arise during intense meditation. This may reflect an increase of visual resources allocated to process occurring visual stimuli. This finding is in line with the evidence of an increase of higher visual areas involvement in expert meditators in both brain activity (Berkovich-Ohana et al., 2016; Xu et al., 2014) and connectivity (Hasenkamp and Barsalou, 2012; Tang et al., 2017). The reduced expertise-modulated connectivity in nodes within the posterior salience network suggests that salient stimuli may be dropped in favor of a narrow focus of awareness on a single object (Vago and Zeidan, 2016).

In our analysis, none of the connections including the prefrontal cortex was selected when predicting experience, even though this area is employed in several tasks related to FA meditation (Ridderinkhof et al., 2004). This may be related to the optimized performances in sustaining and selective attention tasks associated with long-term practitioners (Lutz et al., 2008). Indeed, with more practice the need for these processes is greatly reduced, described by long-term Buddhist monks as *effortless concentration* (Lutz et al., 2008).

In the influential model proposed by Vago and Zeidan (Vago and Zeidan, 2016), the fronto-parietal network (FPN) and the ventral attention network (VAN) play a key role in coordinating awareness and monitoring of experience in the present moment during OM meditation (Raffone and Srinivasan, 2009). In accordance with this model, we found that the dorsomedial prefrontal cortex, which is a core region of the FPN, shows a high correlation with the left Crus, a node implicated in emotional processing (Stoodley et al., 2012). This connection can play a key role for conscious access to emotional contents during OM meditation. Also, the involvement of nodes of the right executive-control network (RECN) in OM meditation is consistent with the model of Vago and Zeidan (Vago and Zeidan, 2016).

The prediction of expertise during OM meditation indicated the right caudate (RCau), as the most implicated node (Figure 3b). This evidence is in line with the study of Gard (Gard et al., 2015) showing greater widespread functional connectivity of the right caudate with several brain regions as in expert meditators as compared to non-expert meditators. Our finding further highlights the specific involvement of RCau in the OM facet of meditation, which is characterized by a wider conscious access to the fields of experience. Specifically, we found an increased expertise-modulated connection of RCau with nodes in the Language Network, the left angular gyrus (LAngG) and left middle temporal gyrus (LMTG) which can be associated to an enhanced hub function of the caudate in multiple cognitive abilities (Graff-Radford et al., 2017). This evidence is also consistent with an earlier finding by our research group, with the same set of long-term meditators, where the activation of the left parieto-temporal areas in OM meditation have been associated to conscious access and meta-awareness (Manna et al., 2010).

Here, we also found a negative connection weight between the right superior frontal gyrus (RSFG) and the right premotor cortex. This finding can be explained in terms of a reduced enactment (plausibly associated with the preparation of movement) of mental states (and in particular negative emotions) in OM meditation with increased meditation expertise (Lutz et al., 2008). Moreover, again in line with our earlier findings (Manna et al., 2010), OM meditation is suggested to involve mainly left fronto-temporo-parietal cortical regions.

Finally, we found a high correlation of connectivity between left and right hippocampus with meditation expertise, in line with the evidence of hippocampal involvement in meditation (Hölzel et al., 2011; Tang et al., 2015). Given that, the hippocampus is a key convergence zone in the brain integrating episodic information (Eichenbaum, 2004), an increased expertise-modulated connectivity between the left and right hippocampus might underlie an enhanced conscious access to episodic information and related emotional contents (Lutz et al., 2008), as well as an enhanced process of interoception (Dudley and Stevenson, 2016), two key aspects of OM meditation.

OM meditation is often practiced after FA meditation (Lutz et al., 2008), since both meditation forms involve a narrow focus on the present moment experience and a detachment from reactivity and judgment patterns (Vago and Zeidan, 2016). Therefore, a shared pattern of connectivity that is predictive of the meditation expertise can be expected.

Here the model revealed a high implication of the posterior insula, which is characterized by decreased connections with the right thalamus and the left angular gyrus. Posterior insula can indeed be involved in both FA and OM forms of meditation given its role in body and present moment-awareness, including breath sensations (Kuehn et al., 2016; Marchand, 2014). Moreover, a reduced functional coupling of posterior insula with the right thalamus and associative areas such as the left angular gyrus might prevent broadcasting of thalamo-cortico-limbic signals associated to emotional reactivity with a potential interference during both FA and OM meditation forms (see also (Raffone and Srinivasan, 2009)). This evidence appears also to converge with our previous study revealing a negative correlation between activation of posterior insula and meditation expertise (Manna et al., 2010).

We demonstrated that the age of meditators can be accurately predicted using fcMRI. Specifically, the analysis revealed that the caudate-thalamus connections within the Basal Ganglia network have the highest importance for the prediction. Indeed, it has been found that such regions decrease their thickness with ageing (Bauer et al., 2015). Moreover, it has been shown that the caudate nucleus has a great importance in practitioners of yoga and insight meditation (Gard et al., 2015) and that lesions of this region have a profound effect on several behavioral and cognitive abilities (Graff-Radford et al., 2017).

Our findings also revealed that thalamic connections are central for the prediction of age. This result appear to be consistent with the ageing effects on the white matter thalamic microstructure observed in long-term meditators (Laneri et al., 2016). A recent study found an increased connectivity of thalamic regions correlated with better performance in a group of older vs. young participants performing a spatial-memory task (Goldstone et al., 2018), our results appears to support the evidence of the beneficial effects of meditation on neurocognitive decline in ageing (Farb et al., 2013; Gard et al., 2015; Luders et al., 2016).

We also found an involvement of insular regions in age prediction; this findings seems related to studies reporting greater fractional anisotropy of white matter in both right and left insula in long-term meditators (Laneri et al., 2016).

Finally, it has to be noted that our experiment was not designed to elucidate the beneficial effects of meditation on ageing, since we were only predicting age from functional connectivity during meditation. However, the stability of our pattern despite the meditation style suggest a persistent change of age-specific modulation of functional connectivity that may reflect a reduction of neurocognitive decline with ageing (Farb et al., 2013; Pickut et al., 2013).

Our study showed that fMRI connectivity patterns within and between multiple key brain networks can differentially predict meditation expertise and age of long-term meditators, with different patterns in focused attention and open monitoring meditation forms. The patterns of connectivity that predicted meditation expertise depended on the meditation style, while connectivity patterns that predicted age were the same for both meditation styles. These findings suggest that brain connectivity patterns within and between brain networks are differentially shaped by meditation expertise and age, with differential influences of focused attention and open monitoring meditation forms.

According to our hypotheses, we found that the FA meditation connectivity pattern modulated by meditation expertise includes nodes and connections implicated in focusing, sustaining and monitoring attention, such as ACC, IPS and AngG. The connectivity pattern during OM meditation includes nodes associated with cognitive and affective monitoring and control, such as dorsomedial prefrontal cortex and the left Crus, as well as with interoception and emotional awareness such as hippocampus and the right caudate. Our findings also highlight the role of posterior insula as a key region in both FA and OM meditation forms. In addition, our results suggest that the caudate may have a central role in the beneficial effects of meditation on cognitive decline and aging.

Possible limitations of our study are the relative small sample size used (Cui and Gong, 2018), that is a common trait of several meditation studies (Tang et al., 2015) and the lack of objective methods to confirm the good execution of the tasks. However, the robustness of the statistical analysis used suggests that the conclusions are not over-inflated. In addition, the dedicated practice of the involved Theravada Buddhist monks in both FA and OM meditation forms might have compensated for these limitations.

In conclusion, our study highlights a long-term effect of meditation practice on multivariate patterns of functional connectivity, in a group of expert meditators, and suggests that meditation expertise is associated with specific neuroplastic changes in connectivity patterns within and between multiple brain networks, with differential effects of FA and OM meditation practices on such patterns.

In particular, while the connectivity pattern associated with FA meditation includes nodes and connections implicated in focusing, sustaining, and monitoring attention the connectivity pattern associated with OM meditation includes nodes and connections associated with cognitive and affective monitoring and control, as well as with interoception. Thus, the two form of meditation may differentially contribute to counteract the effects of neurocognitive decline with ageing by neuroplasticity within and between brain networks.

## Acknowledgements

We would like to thank the monks of Santacittarama Monastery for their outstanding participation in our study. We would also like to thank Carlo Sestieri, Simone Di Plinio, Antea D’Andrea and Annalisa Tosoni for useful discussions.

## Funding

Dr. Antonino Raffone has been supported by a grant from the BIAL Foundation (Portugal) on the project “Aware Mind-Brain: bridging insights on the mechanisms and neural substrates of human awareness and meditation”. This work was supported by the "Departments of Excellence 2018-2022" initiative of the Italian Ministry of Education, University and Research for the Department of Neuroscience, Imaging and Clinical Sciences (DNISC) of the University of Chieti-Pescara.

